# Astrocyte mitochondria produce nitric oxide from nitrite to modulate cerebral blood flow during brain hypoxia

**DOI:** 10.1101/2022.03.13.484128

**Authors:** Isabel N Christie, Shefeeq M Theparambil, Maxim Doronin, Patrick S Hosford, Alexey Brazhe, Adrian Hobbs, Alexey Semyanov, Andrey Y Abramov, Plamena Angelova, Alexander V Gourine

## Abstract

In low oxygen conditions, increases in cerebral blood flow maintain brain oxygen delivery but the mechanisms underlying hypoxia-induced dilations of cerebral vasculature are incompletely understood. Here we show that astrocytes accumulate nitrite and in response to hypoxia produce nitric oxide via mitochondrial reduction of nitrite by a molybdenum-containing enzyme sulfite oxidase. These data suggest that astrocytes can modulate cerebral blood flow in accord with brain tissue oxygenation via mitochondrial production of nitric oxide.

## Introduction

The brain is extremely vulnerable to reduction in oxygen supply (hypoxia) due to the exceptionally high metabolic rate associated with the activities of billions of neurons processing information^1^. If brain blood supply would suddenly cease, the cerebral oxygen content would be enough to maintain neuronal function for merely a few seconds^2,3^. Specialised peripheral (arterial) oxygen sensors are located outside the CNS (in the carotid and aortic bodies) and are not sensitive to regional brain differences in oxygen supply or local brain tissue hypoxia. This suggests the necessity of an intrinsic brain mechanism that can sense oxygen^4^, and dynamically adjust the brain tissue perfusion in accord with regional differences in oxygen concentration, and/or increase global cerebral blood flow (CBF) during systemic hypoxia.

The cerebral vasculature is sensitive to arterial hypoxemia and/or brain tissue hypoxia, responding with dilations and increases in blood flow to maintain brain oxygen delivery^5,6^. Although several studies have addressed the potential roles of certain K^+^ channels, Ca^2+^ channels, low pH, lactate-, ATP-, adenosine- and NO-mediated signalling^5,6^, the exact mechanisms underlying the cerebrovascular responses to brain hypoxia remain incompletely understood. This is important, as significant gradients of brain tissue partial pressure of oxygen (PO_2_) occur even at normal arterial PO_2_ and saturation^7,8^, suggesting that regions of the brain are often exposed to a low oxygen environment^9,10^.

All penetrating and intraparenchymal cerebral blood vessels are wrapped by the end-feet of astrocytes, - omnipresent multifunctional glial cells that control cerebral vasculature via Ca^2+^-mdependent release of vasoactive signalling molecules^11–13^. In this study we evaluated the potential role of astrocytes in the mechanisms underlying hypoxia-induced increases in brain parenchymal blood flow. First, we tested the hypothesis that the direct oxygen sensitivity of astrocytes, which manifests as intracellular Ca^2+^ responses to hypoxia^14^, is responsible for the dilations of cerebral arterioles associated with these astrocytes.

## Results

Using 2-photon imaging in anesthetized and artificially ventilated rats, we recorded robust dilations of cortical arterioles (by 36±3%; 65 vessels recorded in 23 animals) when the concentration of oxygen in the inspired air was lowered to 10% (Figure 1a-c). Reproducible dilations of cortical vessels occurred immediately at the hypoxic stimulus onset (Figure 1b,c). Cortical astrocytes loaded with a Ca^2+^-sensitive dye Oregon Green BAPTA 1 AM responded to systemic hypoxia with increased frequency of Ca^2+^ signals in perivascular end-feet and somas (Figure 1b; Supplementary Figure 1), as reported previously^14^. Hypoxia-induced arteriolar dilations occurred simultaneously with Ca^2+^ signals in astroglial cell bodies, but the latency to the onset of perivascular end-feet Ca^2+^ responses lagged behind vessel dilation onset in most trials (Figure 1b,c). There was also no correlation between the magnitude of the vessel dilation and the Ca^2+^ responses in astrocyte somas and end-feet (Figure 1d), suggesting that cerebrovascular responses to hypoxia are not mediated by Ca^2+^-dependent release of vasoactive substances by astrocytes.

**Figure 1.**
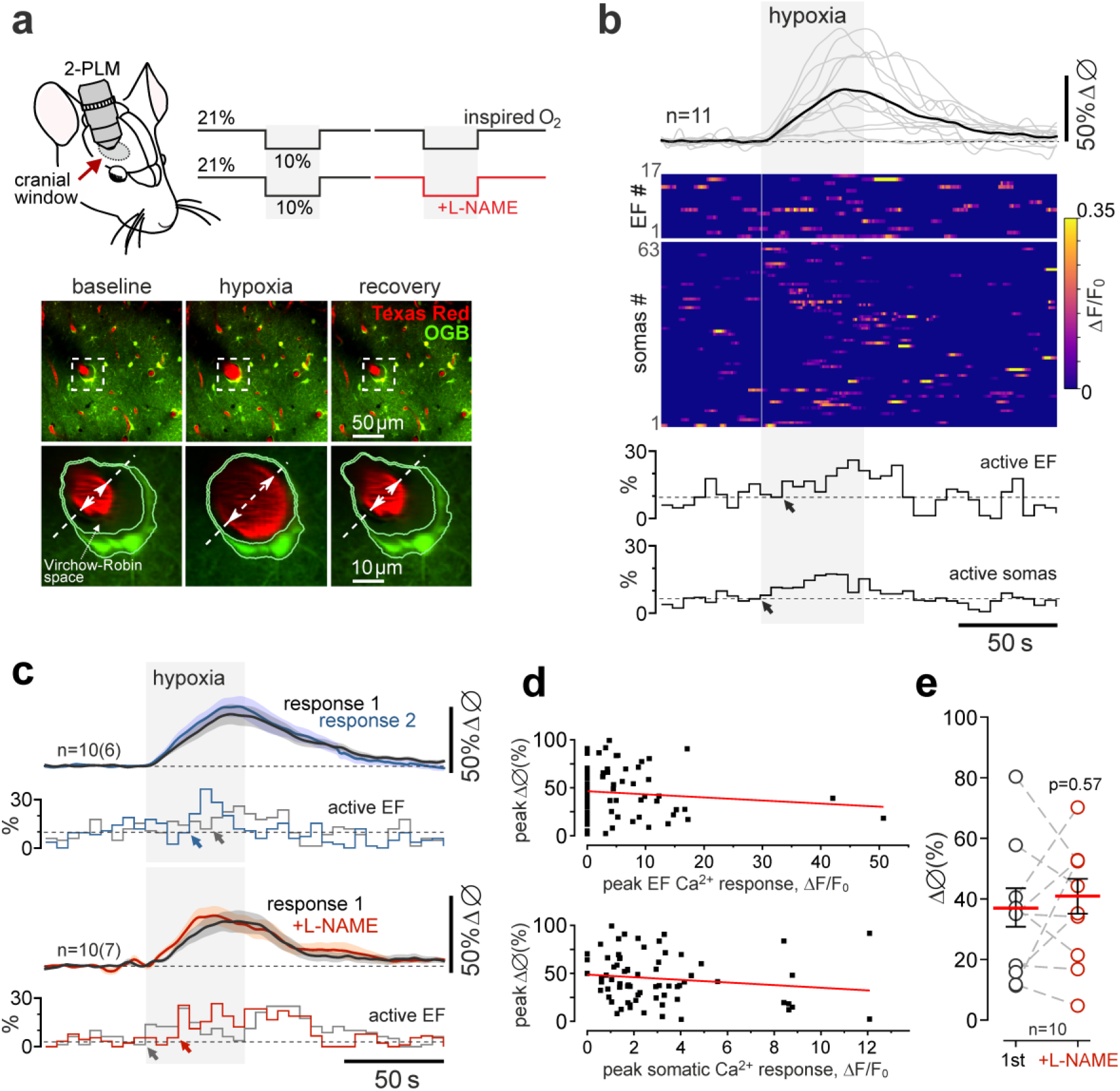
Hypoxia-induced vascular responses in the cerebral cortex. (**a**) Diagram of the experimental protocol in anesthetized rats instrumented for two-photon imaging of cortical blood vessels (visualized with intravascular dye Texas Red) and Ca^2+^ in cortical astrocytes (using a Ca^2+^-sensitive dye Oregon Green BAPTA 1 AM; OGB). Representative images were taken near the cortical surface at baseline, at the peak of the response induced by 10% inspired O_2_ and after a complete recovery from hypoxia. (**b**) Hypoxia-induced changes in the diameter of penetrating cortical arterioles (11 vessels recorded in 6 animals), Ca^2+^ signals in perivascular endfeet (EF; 17 regions of interest; ROIs) and cell bodies (63 ROIs) of cortical astrocytes. Arrows point to the onset of astrocyte Ca^2+^ end-feet (EF) and somatic responses. (**c**) Summary data showing overlaid profiles of changes in the diameter of penetrating cortical arterioles and Ca^2+^ signals in perivascular endfeet in response to two sequential episodes of hypoxia (10% inspired O_2_). Hypoxia triggered reproducible dilations of cortical vessels which were not affected in conditions of established systemic nitric oxide synthase (NOS) blockade with N(ω)-nitro-L-arginine methyl ester (L-NAME; 10 mg kg^−1^). Arrows point to the onset of astrocyte Ca^2+^ end-feet responses. (**d**) Scatterplots showing relations between the hypoxia-induced Ca^2+^ responses in astrocyte end-feet and cell bodies and the corresponding cerebrovascular responses. (**e**) Summary data illustrating peak hypoxia-induced increases in cortical arteriole diameter in control conditions and following systemic NOS blockade with L-NAME. The data are shown as individual values and means ± SEM. *p* value, Wald test on linear mixed effect model with random intercepts.

There is evidence that hypoxia-induced increases in cerebral blood flow are mediated by nitric oxide (NO)^6^. However, we found that in our experimental conditions hypoxia-induced dilations of cortical arterioles in response to 10% inspired O_2_ were unaffected by systemic nitric oxide synthase (NOS) blockade with N(ω)-nitro-L-arginine methyl ester (L-NAME) (Figure 1c,e; Supplementary Figure 2); these results are consistent with the data obtained in humans^15^. However, NO can also be produced following reduction of the nitrite anion (NO_2_^−^), - an alternative mechanism requiring an electron donor and haem- or molybdenum-containing protein^16–18^.

Astrocytes have a loosely assembled mitochondrial respiratory chain, resulting in poor mitochondrial respiration, and propensity to electron leak^19^. We next tested the hypothesis that in low oxygen conditions these electrons are used to generate NO via reduction of nitrite in astrocytes. We used a fluorescent NO indicator DAR-4AM^20^ and genetically encoded NO sensor geNOp^21^ to record NO production by astrocytes and neurons in culture (Figure 2). Both indicators reported robust hypoxia-induced increases in NO production by astrocytes in response to the displacement of oxygen in the incubation medium with argon (Figure 2a,b,e). Hypoxia-induced NO production in astrocytes was unaffected by inhibition of NOS with L-NAME (100 μM) (Figure 2c,e) or in the presence of a mitochondrial antioxidant MitoQ (500 nM; MitoQ scavenges reactive oxygen species, but not nitric oxide) (Figure 2c,e). Hypoxia had no effect on NO production in cultured cerebellar neurons (known to express high level of neuronal NOS) (Figure 2d,e). Experiments conducted in organotypic cortical slices, involving monitoring of PO_2_ in the medium, showed that astrocytes start to produce NO when PO_2_ falls below the threshold of 17±2 mmHg (Figure 2f).

**Figure 2.**
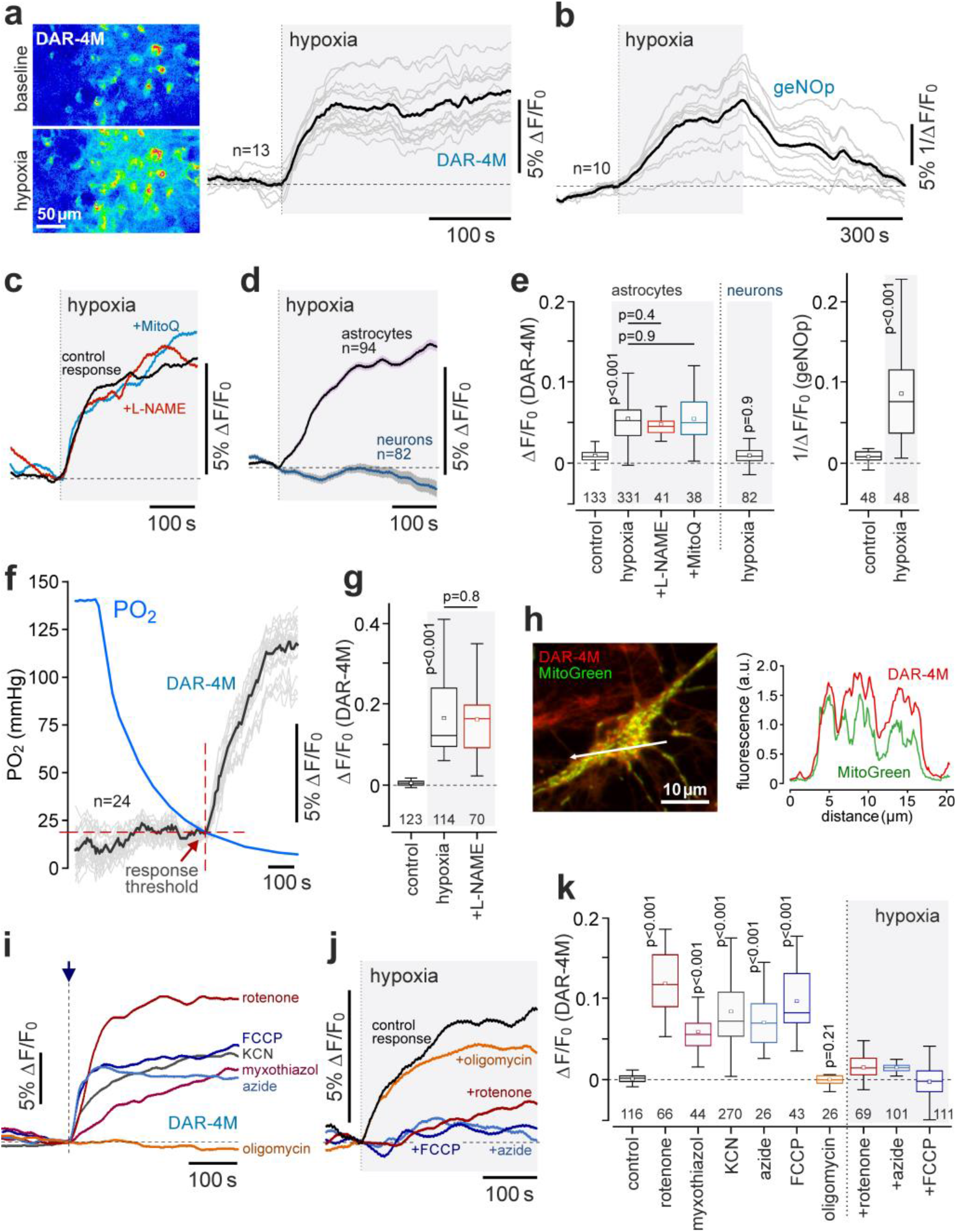
Astrocytes produce nitric oxide in response to hypoxia. (**a**) Hypoxia-induced changes in fluorescence of a nitric oxide (NO)-sensitive dye DAR-4M in cultured astrocytes. Representative images were taken at baseline and at the peak of the response to hypoxia induced by the displacement of oxygen with argon. Traces illustrate representative examples of individual (grey) and averaged (black) changes in DAR-4M fluorescence recorded in 13 astrocytes in one experiment. (**b**) Hypoxia-induced changes in fluorescence of a genetically-encoded NO sensor geNOp expressed in cultured astrocytes. Traces depict individual (grey) and averaged (black) changes in geNOp fluorescence recorded in 10 astrocytes in one experiment. (**c**) Representative profiles of hypoxia-induced NO production by astrocytes in the absence and presence of NOS inhibitor L-NAME (100 μM) and ROS scavenger MitoQ (500 μM). (**d**) Averaged profiles of hypoxia-induced changes in NO production recorded in cortical astrocytes and cerebellar granule neurons. (**e**) Summary data illustrating peak changes in DAR-4M and geNOp fluorescence (reporting NO production) recorded in cortical astrocytes in response to hypoxia, also showing the effects of L-NAME and MitoQ on astrocyte responses and peak hypoxia-induced neuronal responses. (**f**) Simultaneous recordings of NO production by astrocytes in organotypic slices of the cerebral cortex and partial pressure of oxygen (PO_2_) at the surface of the slice, illustrating the PO_2_ threshold of the astroglial response to hypoxia. (**g**) Peak hypoxia-induced changes in DAR-4M fluorescence recorded in organotypic cortical slices in the absence and presence of L-NAME. (**h**) Labelling of astrocyte mitochondria with MitoTracker Green shows mitochondrial accumulation of DAR-4AM. *Right:* Fluorescence intensity profile following the arrow (white) illustrating co-localization of MitoTracker Green and DAR-4AM. (**i**) The effect of hypoxia on NO production in astrocytes is mimicked by inhibition of the mitochondrial electron transport chain or mitochondrial uncoupling (rotenone, 2 μM; myxothiazol, 3 μM; KCN, 100 μM; azide, 1 mM; FCCP, 1 μM; oligomycin, 2 μM). (**j**) Inhibition of the mitochondrial electron transport chain (rotenone, azide) or mitochondrial uncoupling (FCCP) occlude the effect of hypoxia on NO production in astrocytes. (**k**) Peak changes in DAR-4M fluorescence recorded in cortical astrocytes in response to inhibition of the mitochondrial electron transport chain or mitochondrial uncoupling, and in response to hypoxia in conditions of mitochondrial electron transport chain inhibition or mitochondrial uncoupling. In **c**, **i** and **j** each trace shows averaged changes in DAR-4M fluorescence recorded in 8-15 individual astrocytes. *p* values, ANOVA.

Labelling of astrocyte mitochondria with MitoTracker Green showed a preferential mitochondrial accumulation of DAR-4AM indicator (Figure 2h), suggesting that the mitochondrion is the potential site of NO generation during hypoxia. Next we found that the effect of hypoxia on NO production in astrocytes could be mimicked by chemical inhibition of the mitochondrial electron transport chain (chemical hypoxia). Inhibition of mitochondrial complex I with rotenone (2 μM), inhibition of complex III with myxothiazol (3 μM), inhibition of complex IV with cyanide (KCN, 500 μM) or azide (1 mM) or the mitochondrial uncoupling with FCCP (1 μM) facilitated NO generation by astrocytes and blocked the effect of hypoxia on NO production in these cells (Figure 2i-k). Blockade of complex V with oligomycin (2 μM) had no effect on hypoxia-induced NO production by astrocytes (Figure 2i-k). Astrocyte NO production induced by chemical hypoxia (KCN, myxothiazol) was unaffected by L-NAME (Supplementary Figure 3a,b). Inhibition of mitochondrial respiration with rotenone (2 μM) or KCN (500 μM) had no effect on NO production in cultured cerebellar and cortical neurons (Supplementary Figure 4).

Measurements of [NO_2_^−^] in astrocytes or isolated astrocyte mitochondria following their incubation with NaNO_2_ (0, 5, 10, 30, 100 μM) showed that in aerobic conditions astrocyte mitochondria accumulate nitrite (Figure 3a). In the presence of supplemental NO_2_^−^ (100 μM) (but not 100 μM nitrate, NO_3_^−^), NO production by astrocytes in response to hypoxic or chemical hypoxia (KCN) was markedly enhanced (Figure 3b,c). This effect was blocked by inhibition of anion transport with 4,4’-Diisothiocyano-2,2’-stilbenedisulfonic acid (100 μM) (Supplementary Figure 3c), suggesting the existence of mechanism(s) that transport nitrite across the cell and mitochondrial membranes.

**Figure 3.**
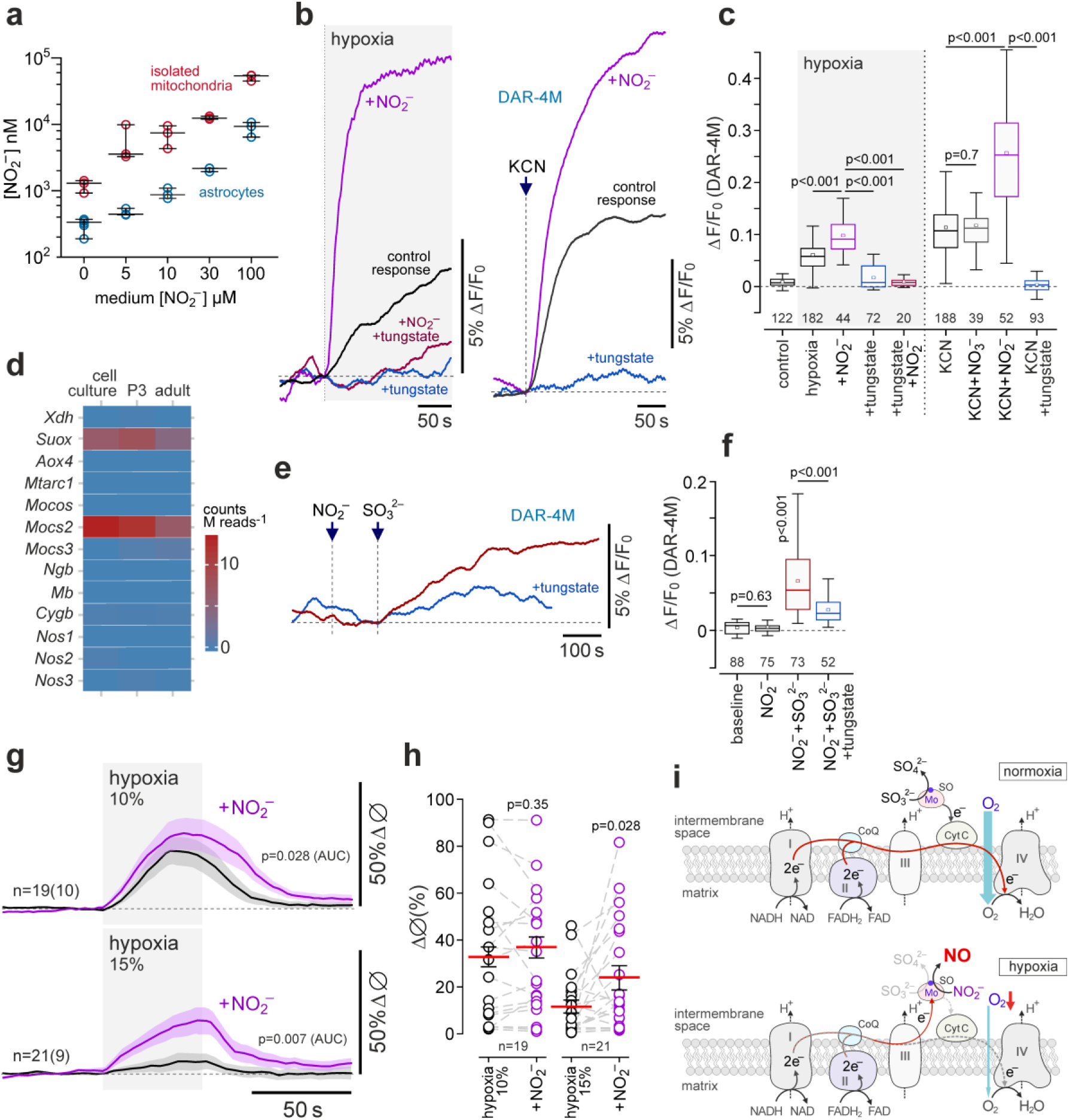
During hypoxia astrocytes produce nitric oxide from nitrite by a molybdenum-containing enzyme sulfite oxidase. (**a**) Astrocyte mitochondria accumulate nitrite. Summary data illustrating measurements of nitrite concentration ([NO_2_^−^]) in isolated astrocyte mitochondria and whole astrocytes that were incubated for 1 h with sodium nitrite (0-100 μM) in aerobic conditions. (**b**) Astrocyte NO production induced by hypoxia or inhibition of complex IV with KCN (100 μM) is augmented by supplemental nitrite (100 μM) and is abolished in cells treated with tungstate (0.5 mM), which replaces molybdenum with tungsten in molybdopterin-containing enzymes. (**c**) Summary data illustrating peak changes in DAR-4M fluorescence (reporting NO production) recorded in cortical astrocytes in response to hypoxia or mitochondrial inhibition with KCN in the absence or presence of supplemental nitrate (NO_3_^−^), nitrite and after the treatment with tungstate. *p* values, ANOVA. (**d**) RNAseq data illustrating the expression of genes encoding metalloproteins with nitrite reductase activity and other relevant proteins in cultured cortical astrocytes, and acutely isolated cortical astrocytes of neonatal (P3) and young adult rats. *Xdh*, xanthine dehydrogenase/oxidase; *Suox*, sulphite oxidase; *Aox4*, aldehyde oxidase 4; *Mtarc1*, mitochondrial amidoxime reducing component; *Mocos*, molybdenum cofactor sulfurase; *Mocs2* and *3*, molybdenum cofactor synthesis genes; *Ngb*, neuroglobin; *Mb*, myoglobin; *Cygb*, cytoglobin; *Nos1*, neuronal NOS; *Nos2*, inducible NOS; *Nos3*, endothelial NOS. (**e**) Sulfite (SO_3_^2-^; 0.5 mM) triggers NO production by astrocytes in the presence of nitrite (100 μM). (**f**) Peak changes in DAR-4M fluorescence recorded in cortical astrocytes in response to nitrite, sulfite in the presence of supplemental nitrite, and in response to sulfite after the treatment of cells with tungstate. *p* values, Kruskal-Wallis ANOVA. (**g**) Hypoxia-induced vascular responses in the cerebral cortex are augmented following systemic treatment with nitrite (1 mg kg^−1^, i.v.). Shown are overlaid profiles of changes in the diameter of penetrating cortical arterioles in response to two sequential episodes of hypoxia (10% or 15% inspired O_2_) before and after administration of nitrite. (**h**) Summary data illustrating peak hypoxia-induced increases in cortical arteriole diameter in control conditions and after nitrite administration. The data are shown as individual values and means ± SEM. *p* values, Wald test on linear mixed effect model with random intercepts. (**i**) Schematic drawing of the mitochondrial electron transport chain illustrating the proposed mechanism of NO production by astrocytes in response to hypoxia. In aerobic (normoxic) conditions sulfite oxidase (SO) oxidases sulfite to sulfate and transfers electrons to cytochrome *c*. In low oxygen conditions when mitochondrial respiration is inhibited and cytochrome *c* is fully reduced, sulfite oxidase transfers an electron and facilitates proton donation to reduce nitrite to NO.

Several metalloproteins can facilitate generation of NO by transferring electrons for nitrite reduction^16,17^. To identify the astroglial enzyme responsible for NO_2_^−^ reduction in hypoxic conditions, we performed RNAseq to evaluate the expression of haem-containing and molybdenum (Mo)-bound molybdopterin cofactor proteins and related enzymes in cultured astrocytes, as well as in astrocytes acutely isolated from the cerebral cortex of neonatal (p3) and adult rats (Figure 3d). Amongst the metalloproteins capable of catalysing nitrite reduction and related factors, high expression of genes encoding Mo-containing mitochondrial enzyme sulfite oxidase (*Suox* gene) and molybdenum cofactor synthesis protein (*Mocs2* gene; involved in the synthesis of molybdopterin cofactor, required for biochemical activation of molybdenum) was detected in cortical astrocytes (Figure 3d). Analysis of data from a publicly available mouse brain transcriptome database^22^ confirmed high expression of *Suox* and *Mocs2* in cortical astrocytes (Supplementary Figure 5).

In the presence of tungstate, tungsten effectively replaces molybdenum in molybdopterin cofactor-containing proteins^23^. We found that after 24 h incubation with sodium tungstate (0.5 mM), cultured astrocytes were no longer able to generate NO in response to hypoxic or chemical hypoxia (Figure 3b,c), indicating that a molybdopterin cofactor protein is responsible for nitrite reduction in these conditions. Xanthine oxidase and aldehyde oxidase (both cytosolic proteins) do not appear to be involved as potent inhibitors of these enzyme, allopurinol (50 μM) and oxypurinol (20 μM)^16^, had no effect on hypoxia-induced NO production by astrocytes (Supplementary Figure 3d). Another molybdenum-containing enzyme - sulfite oxidase - catalyzes the oxidation of sulfite (SO_3_^2-^) in the intermembrane space of the mitochondria^24^. We next hypothesised that if sulfite oxidase is responsible for nitrite reduction in astrocytes, then substrate (sulfite) supplementation should trigger NO production in these cells. Indeed, in the presence of 100 μM NO_2_^−^, addition of sulfite (SO_3_^2-^; 0.5 mM) triggered robust NO production by astrocytes in aerobic conditions (Figure 3e,f). NO production induced by SO_3_^2-^ was markedly reduced in astrocytes treated with tungsten (Figure 3e,f). These data show that in astrocyte mitochondria sulfite oxidase can effectively transfer electrons for nitrite reduction.

We next determined whether the identified mechanism of NO production contributes to the hypoxia-induced increases in cerebral blood flow. If this hypothesis is correct, then supplemental NO_2_^−^ would be expected to augment the dilations of cerebral vasculature in response to decreases in the level of inspired O_2_. Systemic administration of NO_2_^−^ (1 mg kg^−1^) had no effect on the amplitude of cortical arteriole dilations induced by 10% inspired O_2_ but increased the overall response (11.9±0.3 AUC *vs* 11.2±0.2 AUC in control conditions; 19 vessels recorded in 10 animals; p=0.028; Figure 3g,h). This effect of supplemental NO_2_^−^ was relatively small and could be potentially explained by 10% inspired O_2_ causing near-maximal arteriolar dilations and exhaustion of the cerebrovascular reserve. Therefore, we next determined the effect of supplemental NO_2_^−^ on cortical arteriole responses to moderate hypoxia induced by 15% inspired O_2_ (Figure 3g,h). Peak dilations of cortical arterioles induced by 15% inspired O_2_ were increased by 100% following systemic administration of NO_2_^−^ (diameter increase by 24±5% *vs* 12±3% in control conditions; 21 vessels recorded in 9 animals; p=0.028; Figure 3g,h). Nitrite administration had no effect on Ca^2+^ responses in astrocyte cell bodies and perivascular endfeet induced by 10% or 15% inspired O_2_ (Supplementary Figure 1b).

## Discussion

Together these data suggest that astrocytes accumulate NO_2_^−^ and during periods of hypoxia generate NO by mitochondrial reduction of NO_2_^−^ to dilate neighbouring cerebral blood vessels and increase brain blood flow. That the effect of chemical inhibition of mitochondrial respiration mimics, but also occludes the effect of hypoxia, on NO production by astrocytes, points to mitochondria as the site of NOS-independent NO generation by these cells. Blockade of NO production by tungsten indicates that in astrocytes a mitochondrial molybdenum-molybdopterin containing enzyme is responsible for hypoxia-induced reduction of nitrite. Animals express only four members of this family of enzymes^16,17^ and our transcriptome analysis demonstrates high relative (compared to other known enzymes and metalloproteins, capable of producing NO) expression of sulfite oxidase in cortical astrocytes. In aerobic conditions and in the presence of supplemental NO_2_^−^, astrocytes generate NO in response to SO_3_^2-^. Collectively, these lines of evidence indicate that sulfite oxidase is the key mitochondrial enzyme responsible for NO production by astrocytes in low oxygen conditions (Figure 3i).

In contrast to NO synthesis by the enzymes of the NOS family, generation of vasoactive NO by this mechanism does not require molecular oxygen. This appears to be particularly advantageous in conditions of reduced oxygen supply when rapid signalling by NO is required to increase tissue perfusion. The reduction of nitrite to NO by metalloproteins that transfer an electron and facilitate proton donation is favoured by acidic pH^16,17^. Being structurally similar to plant nitrate reductase^25^, sulfite oxidase is located in the proximity to cytochrome *c* within the mitochondrial intermembrane space^26^ where the conditions for NO_2_^−^ reduction appear to be ideal (Figure 3i), especially in astrocytes as these cells have loosely assembled mitochondrial respiratory chain and propensity to electron leak^19^. We hypothesise that the identified mechanism of NO production by astrocytes dynamically matches regional cerebral perfusion with brain tissue oxygenation and contributes to increases in global cerebral blood flow that occur during systemic hypoxia.

## Methods

All animal experimentations were performed in accordance with the European Commission Directive 2010/63/EU (European Convention for the Protection of Vertebrate Animals used for Experimental and Other Scientific Purposes) and the UK Home Office (Scientific Procedures) Act (1986) with project approval from the Institutional Animal Care and Use Committee of the University College London. The animals were group-housed and maintained on a 12-h light/dark cycle (lights on 07:00) and had *ad libitum* access to water and food. The rats were housed in a temperature-controlled room at 22°C with 55±10% relative humidity.

### *Two-photon imaging* in vivo

Young Sprague-Dawley rats (100–150 g) were used to record hypoxia-induced cortical vascular responses and Ca^2+^ signals in cortical astrocytes. The animal was anesthetised with isoflurane (2-4% in O_2_-enriched air). The femoral artery and vein were cannulated for the recordings of the arterial blood pressure and administration of drugs, respectively. Isoflurane was discontinued after intravenous delivery of urethane (1 g kg^−1^) and α-chloralose (50 mg kg^−1^). Adequate anaesthesia was ensured by maintaining stable levels of the arterial blood pressure and heart rate showing lack of responses to a paw pinch. The trachea was cannulated and the animal was mechanically ventilated using a positive pressure ventilator (tidal volume: ~1 ml per 100 g of body weight; frequency: ~60 strokes min^−1^) with oxygen enriched air (~30% O_2_). The head of the animal was secured in a stereotaxic frame and a small circular craniotomy (~3 mm^2^) was made in the parietal bone above the somatosensory cortex.

Cortical cells were loaded with a Ca^2+^-sensitive dye Oregon Green BAPTA 1 AM (OGB). OGB was first dissolved in DMSO and Pluronic F127 (20%). The solution containing OGB (1 mM) in artificial cerebrospinal fluid (aCSF; 124 mM NaCl, 3 mM KCl, 2 mM CaCl_2_, 26 mM NaHCO_3_, 1.25 mM NaH_2_PO_4_, 1 mM MgSO_4_, 10 mM D-glucose saturated with 95% O_2_/5% CO_2_, pH 7.4) was delivered by microinjection via a glass micropipette at 2-4 sites within the targeted area of the cortex. The exposed surface of the brain was then covered with agarose (1%) and protected with a glass coverslip secured to the skull with a headplate and acrylic dental cement. An intravascular fluorescent dye Texas Red™ (MW 70,000; Sigma) was administered intravenously. During imaging the animal was neuro-muscularly blocked with gallamine triethiodide (induction: 50 mg kg^−1^, i.v.; maintenance: 10 mg kg^−1^ h^−1^, i.v.). Arterial *P*O_2_, *P*CO_2_, and pH were measured regularly and kept within the physiological ranges by adjusting the tidal volume and the ventilator frequency. The body temperature was maintained at 37.0±0.5°C.

Vascular and cellular responses in the cortex were recorded using an Olympus FV1000 microscope (Olympus), equipped with MaiTai HP DeepSee laser (Spectra-Physics). A 25x water-immersion objective (XLPlan N, NA 1.05; Olympus) was used. Fluorophores were excited in two-photon XYZ-t mode at 800 nm and images were acquired between 100-200 μm deep from the cortical surface. Cortical arterioles (penetrating and intraparenchymal) were identified anatomically and by the fluorescence of Texas Red. Astrocytes were identified by their characteristic anatomical features such as endfeet. The laser power was kept to a minimum to reduce photo toxicity. Time lapse sequences of recordings were acquired before and after the systemic administration of test compounds for up to 10 min with a period of hypoxia (10% or 15% O_2_ in the inspired air) lasting ~1 min.

### Processing of two-photon imaging data

Rigid-body shifts in the recorded data due to in-plane motion of the tissue were corrected using red fluorescence channel using an average over the first 20 frames as a template. Prior to further analysis all the data was denoised by a two-step truncated singular value decomposition (SVD) projection algorithm. In short, each colour channel video was split to overlapping spatial windows (“patches”, 8×8 pixels), each window was then serialized to form a 2D matrix with single-pixel signals in rows; these matrices were approximated with truncated SVD. In the second step, temporal singular vectors were collected in larger windows and approximated by a second truncated SVD. Inverse SVD transform provided denoised estimate of the fluorescence dynamics within each patch; estimates from overlapping patches were then averaged.

Cortical vessel diameters were measured as described in detail previously^27,28^. For each vessel in the field of view, a line-scan was manually drawn across the widest part of the vessel and the lumen of the vessel was segmented using Chang-Vese active contours with constrains. Cell bodies of astrocytes were identified by an adaptive thresholding algorithm. Regions of interest (ROIs) corresponding to astrocyte somata were manually selected for the analysis. Average intensity of the OGB fluorescence signal was used for Ca^2+^ signals detection. Event detection in each ROI trace was done as described previously^28,29^. Astrocyte endfeet were identified as bright OGB-stained structures encircling penetrating arterioles. To analyse Ca^2+^ responses in the endfeet, a special algorithm was developed to trace the movements of the endfeet associated with vessel diameter changes. In short, a ridge-enhancing filtering was applied to each frame in a crop around a vessel of interest in the green fluorescence channel. These ridge-enhanced frames were resampled to polar coordinates with the origin at the vessel centre, and the endfoot was segmented as a brightness-weighted path from φ=0 to φ=360 within the graph of connected local maxima locations with constraint on path length in each frame. Non-filtered OGB fluorescence signals were then sampled along the circumference of the endfeet in each frame. This kymograph of OGB signal was used for the detection of Ca^2+^ signals in the perivascular endfeet.

### Cell cultures and organotypic slice cultures

Primary cultures of cortical astrocytes were prepared from the brains of rat pups (P2-3 of either sex) as described^30–33^. The animals were euthanized by isoflurane overdose, the brains were removed, and the cortical regions were separated by dissection. After isolation, the cells were plated on poly-D-lysine-coated coverslips and maintained at 37°C in a humidified atmosphere of 5% CO_2_ and 95% air for a minimum of 10 days before the experiments. Viral vector to express genetically encoded NO sensor geNOp (AAV5-CMV-geNOp, titre 1.3×10^11^) was added to the incubation medium after 5-7 days from the time of cell culture preparation.

Neuronal cultures were prepared from the cerebellum as the expression of the neuronal NOS is the highest in this area of the brain^34^. Rat pups (P6-8 of either sex) were euthanized by isoflurane overdose, the brains were removed, and the cerebellum was separated in ice cold Hank’s Balanced Salt Solution (HBSS) buffer. Cells were dissociated after tissue incubation with TrypLE enzyme (Thermo Fisher Scientific; 15 min at 37°C) and suspended in Neurobasal medium containing B-27 supplement, 2 mM L-Glutamine, 25 mM K^+^, 100 U ml^−1^ penicillin, and 0.1 mg ml^−1^ streptomycin. Cells were then plated on poly-D-lysine-coated coverslips (15 mm) and maintained at 37°C in a humidified atmosphere of 5% CO_2_ and 95% air for a minimum of 5 days before the experiments.

Organotypic cortical slice cultures were prepared from the brains of rat pups (P8-10 of either sex), as described in detail previously^14^. The animals were euthanized by isoflurane, the brains were removed and placed in ice-cold HBSS without Ca^2+^, with added 20 mM glucose (total 25.6 mM), 10 mM MgCl_2_, 1 mM HEPES, 1 mM kynurenic acid, 0.005% phenol red, 100 U ml^−1^ penicillin, and 0.1 mg ml^−1^ streptomycin. A sequence of coronal slices (400 μm) was cut at the level of the somatosensory cortex and plated on Millicell-CM organotypic culture membrane inserts (Merck Millipore). Slices were cultured in a medium containing 50% Optimem-1, 25% foetal bovine serum (FBS), 21.5% HBSS; 25 mM glucose, 100 U ml^−1^ penicillin, and 0.1 mg ml^−1^ streptomycin. After 3 days, the plating medium was removed and DMEM medium containing 10% FBS, 2 mM L-Glutamine, 100 U ml^−1^ penicillin, and 0.1 mg ml^−1^ streptomycin was added and subsequently replaced twice a week. Experiments were performed after 7-10 days of incubation.

### Optical recordings of NO production in astrocytes

Optical recordings of changes in NO production in cultured astrocytes were performed using an inverted epifluorescence Olympus microscope, equipped with a cooled CCD camera (Clara, Andor, Oxford Instruments). The cells were loaded with NO sensitive fluorescent probe DAR-4M-AM (Sigma; 10 μM for 30 min incubation at room temperature) or transduced to express genetically encoded NO sensor geNOp (NGFI; Austria). After incubation with the dye the cultures were washed three times with aCSF prior to the experiment. Recordings were performed in a custom-made flow-through imaging chamber at ~32°C in aCSF saturated with 95% O_2_ / 5% CO_2_ (pH 7.4). The rate of chamber perfusion with aCSF was 4 ml min^−1^. To record changes in cytosolic [NO], DAR-4M or geNOp fluorescence were excited by using a Xenon arch lamp and a Optoscan Monocromator (Cairn Research) at 560/10 and 488/10 nm and the florescence emission was recorded at 590 and 535 nm, respectively.

Hypoxic conditions *in vitro* were induced by displacement of oxygen in the medium with argon. All test drugs were applied ~10 min before the hypoxic challenge. Imaging data were collected and analyzed using Andor iQ3 software (Andor, Belfast, UK). All data presented were obtained from at least 6 separate experiments.

### Measurements of partial pressure of oxygen (PO_2_)

In the experiments in organotypic slices, PO_2_ was recorded using optical fluorescence probes (250 μm tip diameter, OxyLite™ system, Oxford Optronix) placed on the surface of the slice as described^14^.

### Nitrite measurements

Accumulation of NO_2_^−^ in astrocytes and astrocyte mitochondria was determined by chemiluminescence assay, as described^35^. After 12 days in culture, astrocytes were washed with phosphate-buffered saline (PBS) and incubated for 60 min in HBSS containing NaNO_2_ in concentration 0, 5, 10, 30 or 100 μM. Samples were then washed with PBS (3x), treated with trypsin (5 min), and centrifuged (at 240×g, 5 min). The supernatant was removed, and the cell pellets were resuspended in PBS to washout the extracellular nitrite. This procedure was repeated twice. After the last wash, the cells were resuspended in 10 ml ddH_2_O for osmotic lysis and the samples were flash-frozen and stored at −80°C until assayed. In a separate set of experiments, pure astroglial cultures were incubated for 60 min with NaNO_2_ (0, 5, 10, 30 or 100 μM), as described above. After harvesting by trypsinization, the cells were washed twice in PBS, incubated on ice for 15 min, and mitochondria were isolated by a series of centrifugations at 600×g, 900×g and 12 000×g. Final (mitochondrial) pellet was then resuspended in ddH_2_O to induce lysis of mitochondria and the samples were stored at −80°C until assayed.

### Isolation and purification of astrocytes for RNA sequencing

Young adult male rats (~100g) and rat pups (P3 of either sex) were used to isolate cortical astrocytes as described in detail previously^31^. The animals were humanely killed by isoflurane overdose and the brains were isolated. The cortex was dissected out and the meninges were removed. The tissue was enzymatically dissociated to make a suspension of individual cells. The samples were passed through a 45 μm Nitex mesh to remove undissociated cell clumps and after addition of myelin removal beads (Miltenyi Biotec), passed through a MACS column (Miltenyi Biotec). The second (positive) magnetic separation was then performed using astrocyte-specific anti-GLAST (ACSA-1) antibodies conjugated to the magnetic beads (Miltenyi Biotec). Anti-O4 selection using magnetic microbeads (Miltenyi Biotec) to separate O4+ immature oligodendrocytes from cell suspensions was also carried out to remove ACSA-1 positive astrocytes contaminated by oligodendrocytes. Cell purity was assessed by anti-GLAST (ACSA-1) phycoerythrin antibody (Miltenyi Biotec) using flow cytometry (CyanADP Cytometer, Beckman Coulter). FACS analysis confirmed >95% purity of isolated astrocytes. Purified cells were harvested by centrifugation at 2000× g for 5 min. The cell pellet was then used for RNA extraction. Separately, cultured astrocytes were individually collected using patch pipettes (tip ~5 μm) made of borosilicate glass. One biological replicate consisted of 20-25 pooled cells from 3-4 cultures.

Total RNA was isolated using the RNeasy Plus Micro Kit (Qiagen) following the manufacturer’s protocol and RNA quality was assessed using the RNA 6000 Pico Kit on a 2100 Bioanalyzer (Agilent Technologies). RNA sequencing, read mapping and expression level estimation were performed as described in detail in^31^. Reads were aligned to the rat reference genome RGSC3.4.64 with TopHat 1.3.3. Cufflinks v1.0.2 was used to assemble and quantitate the transcriptome of each sample. A union set of transcripts in all samples was generated with Cuffcompare, and differential expression was assessed with Cuffdiff. Expression level is reported as fragments per kilobase of transcript sequence per million mapped fragments (FPKM) values.

Expression of genes coding for nitric oxide synthases and all known metalloproteins that possess nitrite reductase activity^16,17^ were compared across the three conditions (cultured cortical astrocytes, and cortical astrocytes acutely isolated from neonatal and young adult rats). Genes included in the analysis were as follows: *Aox4*, aldehyde oxidase 4; *Cygb*; cytoglobin, *Mtarc1*; mitochondrial amidoxime reducing component 1, *Mb*; myoglobin, *Mocos*; molybdenum cofactor sulfurase, *Mocs2*; molybdenum cofactor synthesis 2, *Mocs3*; molybdenum cofactor synthesis 3, *Ngb*; neuroglobin, *Nos1*; nitric oxide synthase neuronal, *Nos2*; nitric oxide synthase inducible, *Nos3*; nitric oxide synthase endothelial, *Suox*; sulfite oxidase, *Xdh*; xanthine dehydrogenase/oxidase, Plots were created using the ggplot2 package in R.

### Statistical analysis

Data were compared by linear mixed-effects model test for nested data with random intercepts, Kruskal-Wallis ANOVA, or one way ANOVA with Tukey post-hoc test, as appropriate. Linear mixed-effects models were used to analyze the effects of L-NAME and NO_2_^−^ on vascular responses and Ca^2+^ signals in astrocyte somata and endfeet; the presence or absence of the drug was treated as a fixed effect, and intercepts for each animal and structure of interest were treated as random effects. *p* values were determined by post-estimation inference using Wald tests. Linear mixed-effect model analysis was performed in “statsmodels” library for Python. The data are shown as individual values and means ± SEM or box-and-whisker plots.

