## Supplemental Information for "Astrocyte mitochondria produce nitric oxide from nitrite to modulate cerebral blood flow during brain hypoxia"

Christie, Theparambil et al.

---

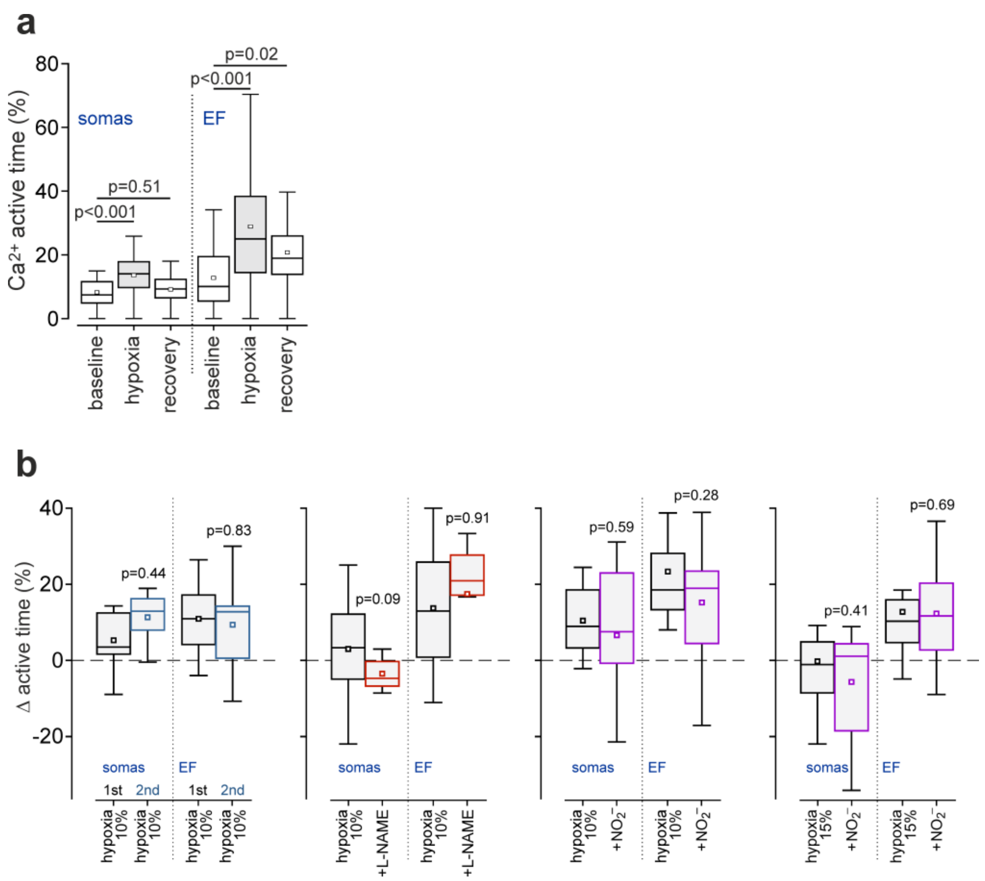


Supplementary Figure 1| (**a**) Summary data illustrating the effect of hypoxia on $Ca^{2+}$ active time (% of time when $[Ca^{2+}]_i$ is elevated) recorded in somas and end-feet of cortical astrocytes. (**b**) Summary data illustrating peak $Ca^{2+}$ responses recorded in somas and end-feet of cortical astrocytes induced by repeated episodes of hypoxia (10% or 15% of $O_2$ in the inspired air) in control conditions, following systemic NOS blockade with L-NAME (10 mg $kg^{-1}$, i.v.), or systemic treatment with nitrite (1 mg $kg^{-1}$, i.v.). In the box-and-whisker plots the central dot indicates the mean, the central line indicates the median, the box limits indicate the upper and lower quartiles and the whiskers show the minimum-maximum range of the data. *p* values, Wald test on linear mixed effect model with random intercepts.

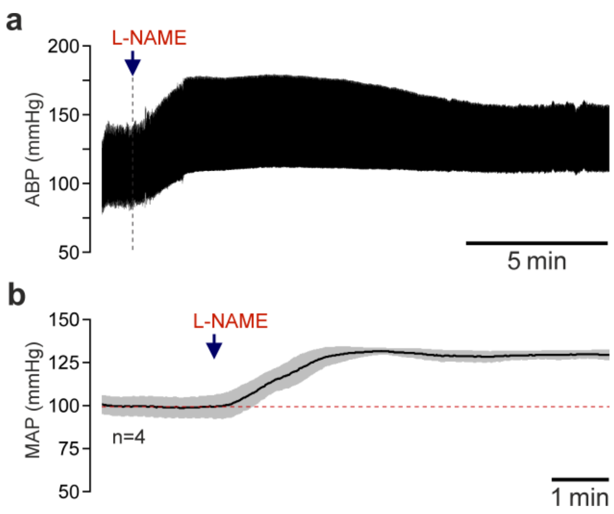


Supplementary Figure 2| (**a**) A representative recording and (**b**) summary data illustrating changes in systemic arterial blood pressure (ABP) after intravenous administration of N(ω)-nitro-L-arginine methyl ester (L-NAME; 10 mg $kg^{-1}$). Sustained elevation of the arterial pressure confirms effective systemic NO synthase inhibition. MAP, mean arterial blood pressure. Data are presented as means ± s.e.m.

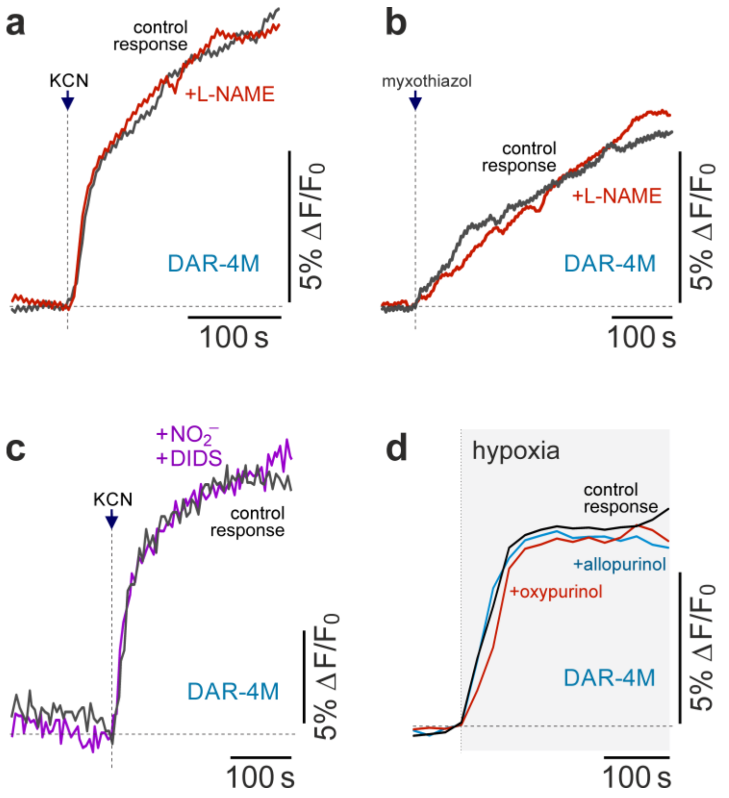


Supplementary Figure 3| (**a**) Representative traces showing changes in fluorescence of a nitric oxide (NO)-sensitive dye DAR-4M in cultured astrocytes in response to mitochondrial complex IV blockade with KCN (500 µM) in the absence and presence of NO synthase (NOS) inhibitor N(ω)-nitro-L-arginine methyl ester (L-NAME; 100 µM). (**b**) Representative traces showing changes in DAR-4M fluorescence in astrocytes in response to mitochondrial complex III blockade with myxothiazol (3 µM) in the absence and presence of L-NAME (100 µM). (**c**) Blockade of anion transport with 4,4'-Diisothiocyano-2,2'-stilbenedisulfonic acid (DIDS; 100 µM) prevents the effect of supplemental nitrite (100 µM) on NO production induced by KCN (500 µM) in astrocytes. (**d**) Representative traces showing hypoxia-induced changes in DAR-4M fluorescence in astrocytes in the absence and presence of xanthine oxidase inhibitors allopurinol (50 µM) or oxypurinol (20 µM).

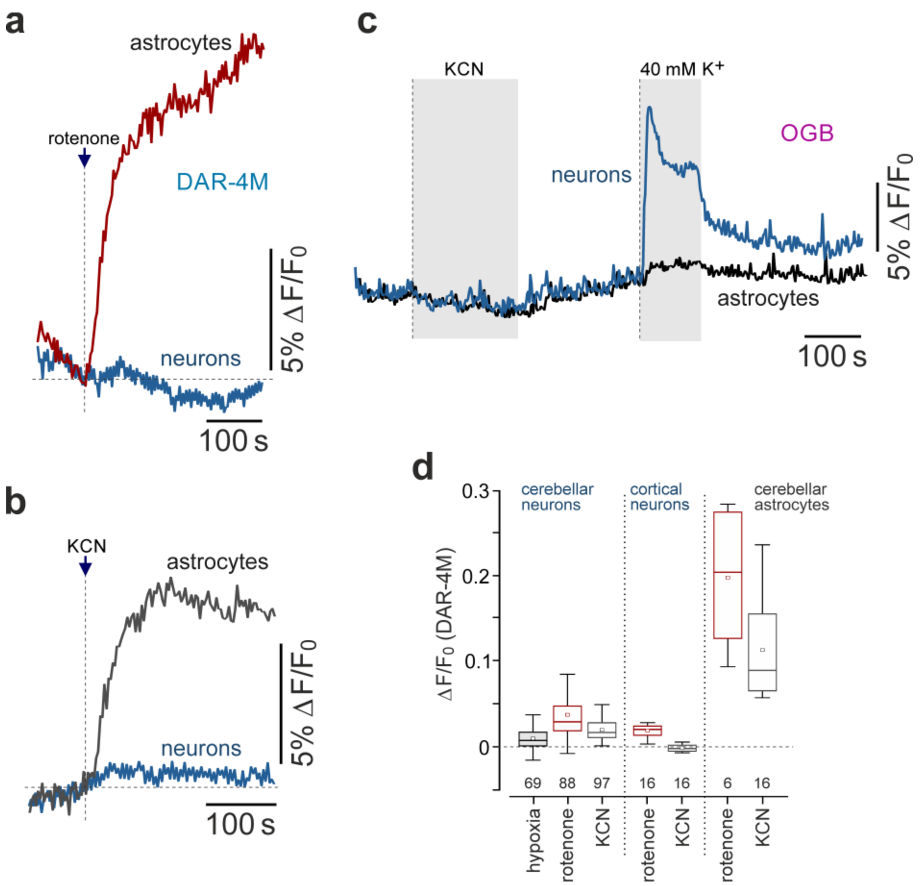


Supplementary Figure 4| (**a**) Representative traces showing changes in fluorescence of NO-sensitive dye DAR-4M in cerebellar granule neurons and astrocytes in response to mitochondrial complex I inhibition with rotenone (2 µM). (**b**) Representative traces showing changes in DAR-4M fluorescence in cerebellar granule neurons and astrocytes in response to mitochondrial complex IV blockade with KCN (500 µM). (**c**) Representative traces showing changes in fluorescence of a $Ca^{2+}$-sensitive dye Oregon Green BAPTA 1 AM (OGB) in cultured cerebellar astrocytes and neurons in response to KCN (500 µM) and KCl (40 mM). Cerebellar granule cells were identified by robust $[Ca^{2+}]_i$ responses to $K^+$. (**d**) Summary data illustrating peak changes in DAR-4M fluorescence recorded in cerebellar granule neurons, cortical neurons, and cerebellar astrocytes in response to inhibition of the mitochondrial electron transport chain with rotenone or KCN.

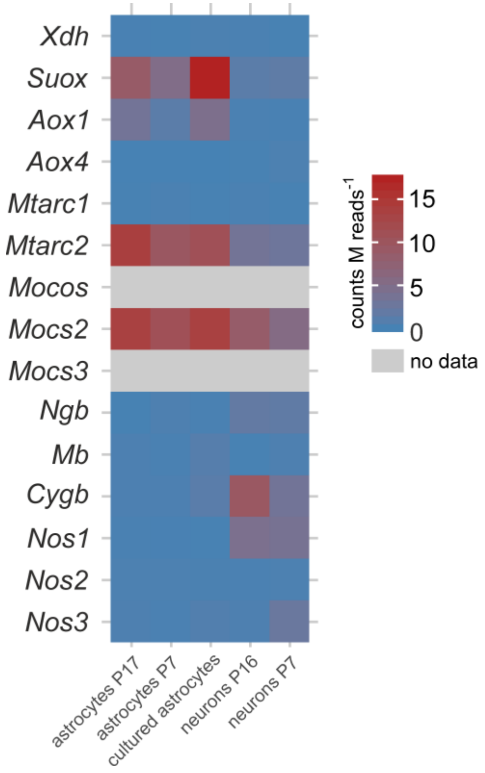


Supplementary Figure 5| Analysis of RNAseq data from a published astrocyte and neuronal transcriptome database (Cahoy et al., 2008) illustrating the relative expression of genes encoding metalloproteins with nitrite reductase activity and other relevant proteins in astrocytes and neurons acutely isolated from the mouse forebrain as well as in cultured astrocytes. The data show that relative expression of the genes of interest is similar to that obtained in cultured and acutely isolated astrocytes of the rat cerebral cortex (Figure 3d). *Xdh*, xanthine dehydrogenase/oxidase; *Suox*, sulphite oxidase; *Aox1*, aldehyde oxidase 1; *Aox4*, aldehyde oxidase 4; *Mtarc1*, mitochondrial amidoxime reducing component 1; *Mtarc2*, mitochondrial amidoxime reducing component 2; *Mocos*, molybdenum cofactor sulfurase; *Mocs2* and *3*, molybdenum cofactor synthesis genes; *Ngb*, neuroglobin; *Mb*, myoglobin; *Cygb*, cytoglobin; *Nos1*, neuronal nitric oxide synthase (NOS); *Nos2*, inducible NOS; *Nos3*, endothelial NOS.
